# Individual genotype but not phenotype predicts river migration success in Atlantic salmon

**DOI:** 10.1101/2023.08.15.553252

**Authors:** Paolo Moccetti, Jonathan D. Bolland, Colin E. Adams, Jessica R. Rodger, Hannele M. Honkanen, Matthew Newton, Angus J. Lothian, Andy D. Nunn, Domino A. Joyce

## Abstract

Migratory species typically undertake demanding long-distance journeys, across different habitat types during which they are exposed to multiple natural and anthropogenic stressors. Mortality during migration is typically high, and may be exacerbated by human-induced pressures. Understanding individual responses to these selection pressures is rarely attempted, because of the challenges of relating individual phenotypic and genetic data to migration success. Here we show distinct Single Nucleotide Polymorphism (SNP) sets significantly differentiated between Atlantic salmon smolts making successful migrations to sea and those that failed to migrate, in two different rivers. In contrast, morphological variation was not diagnostic of migration success. Populations from each river were genetically distinct, and while different genes were possibly implicated in migration success in each river, they related to common biological processes (for example osmoregulation and immune and stress response). Given that migration failure should quickly purge polymorphism at selected SNPs from a population, the question of how genetic diversity in these populations is maintained is an important one. Standing genetic variation could be maintained by different life history strategies and/or environmentally driven balancing selection. Our work highlights the importance of preserving genetic diversity to ensure evolutionary resilience at the population level, and has practical implications for management.

## Introduction

Animal migration has evolved independently many times across the Animal Kingdom (Dingle & Drake, 2007; Bowlin et al., 2010; Shaw, 2016). Migration events can involve large numbers of individuals moving between different habitats and regions, and these events play a key ecological and socio-economic role in natural and human communities (Bauer & Hoye, 2014). Migratory species typically rely on multiple habitats to complete their life cycle and often undertake demanding long-distance journeys exposing themselves to numerous natural and anthropogenic stressors, such as predation, adverse weather conditions, pathogens, pollution, artificial constructions and harvesting (Alerstam, Hedenström & Åkesson, 2003). Mortality during migration is typically high and can be exacerbated by human-induced pressures, such that it impacts upon migratory populations and the ecosystems that depend on them (Wilcove & Wikelski, 2008; Harris et al., 2009; Middleton et al., 2013; Klaassen et al., 2014; Baker et al., 2020). With an ongoing global decline of migratory species (Wilcove & Wikelski, 2008), a better understanding of the factors causing mortality in migration is urgently required to predict responses of migratory populations to future environmental challenges and implement incisive conservation actions.

Recent advances in telemetry technology have made it possible to investigate migratory behaviours of species both temporally and spatially (Doherty et al., 2017; Thorup et al., 2023). This has enabled a better understanding of the exogenous factors directly influencing migration mortality (Thorstad et al., 2013; Hays et al., 2003; Palacín et al., 2017; Weinz et al., 2020). Organisms require a suite of specific morphological, physiological and behavioural adaptive features to successfully complete a migratory cycle (Justen & Delmore, 2022). Given the phenotypic and genetic variation found in most populations, it is reasonable to expect that some genetic or phenotypic traits are more likely to increase migration success than others. However which traits these might be remains poorly understood.

Genomic tools have recently been applied to identify factors regulating migratory behaviour at the population or species level. Several studies have discovered the genetic basis for migratory features such as migration timing and distance, orientation and propensity to migrate, with specific genomic regions linked to these traits (Zhu et al., 2009; Liedvogel, Åkesson & Bensch, 2011; Hecht et al., 2012; O’Malley et al., 2013; Hess et al., 2014; Hecht et al., 2015; Pritchard et al., 2018; Waples, Naish & Primmer, 2020; Justen & Delmore, 2022). However, understanding the genomic basis on which selection could act at an individual level to dictate migration success has rarely been attempted (but see Bourret, Dionne & Bernatchez, 2014), despite the fundamental insights it could provide into how populations might respond to selection, and the implications for management of conservation genetics of migratory species. An important phenotypic trait expected to influence migration success is body morphology (Minias et al., 2013). Morphological variation (i.e. body shape and size) can affect behaviour, resource use, survival and reproductive success of individuals (Wainwright 1994; Skulason & Smith, 1995; Fruciano, Tigano & Ferrito, 2011). The effect of morphology on movement is particularly evident in fish because of a direct link to swimming performance (Pakkasmaa & Piironen, 2000; Fisher & Hogan, 2007; Drinan et al., 2012; Stelkens et al., 2012; Páez & Dodson, 2017). Chapman et al. (2015) found a direct correlation between migration propensity and body shape, while other studies have demonstrated an increased ability and ‘motivation’ to pass river barriers in relation to size, fat content and morphology (Newton et al., 2018; Lothian et al., 2020; Goerig et al., 2020). Nevertheless, research on migration survival and mortality as a consequence of body shape variation (as opposed to size; Kennedy, Gale & Ostrand, 2007; Hostetter et al., 2012; Romer et al., 2013; Furey et al., 2016; Lilly et al., 2022) is still lacking.

Atlantic salmon (*Salmo salar* Linnaeus) is a migratory species of socio-economic importance that has suffered substantial declines over the past 40 years (ICES, 2021) due to multiple abiotic and biotic factors not yet fully understood (Forseth et al., 2017; Dadswell et al., 2022). The Atlantic salmon has a complex life cycle, which includes two long distance migration stages; a long-distance feeding migration from freshwater to sea as a juvenile (smolt) and an adult returning spawning migration from sea to freshwater. In addition it is a philopatric species, accurately homing to its natal spawning grounds (Thorstad et al., 2010). Fidelity to a specific river limits gene flow among populations and has been shown to promote the evolution of local adaption through natural selection, genetic drift, and bottlenecks (Garcia De Leaniz et al., 2007; Fraser et al., 2011). The seaward migration of smolts constitutes a key life-stage for Atlantic salmon and provides an ideal opportunity to study the genetic and phenotypic components that may differentially affect the ability of individual animals to successfully complete their migration. The identification of genetic and phenotypic traits could play a vital role in local management of Atlantic salmon (Bernos, Jeffries & Mandrak, 2020).

Here, we analysed genomic and morphological data of migrating Atlantic salmon smolts in two rivers. We wanted to test to what extent (I) Atlantic salmon populations in the two rivers were genetically distinct, and (II) migration success by seaward migrating smolts could be predicted by specific genomic regions and/or morphological traits.

### Methods

### Sampling, tagging and study design

The study reported here formed part of a wider acoustic telemetry study to examine migratory behaviours and migration success in juvenile salmon (smolts) on their first migration from natal rivers to sea (see Whelan, Roberts & Gray, 2019). Atlantic salmon were captured between 11 April and 3 May 2019 from the rivers Oykel (57°59.640’ N, 4°48.282’ W) and Spey (57°24.960’ N, 3°22.602’ W), Scotland, using 1.5 m diameter rotary screw traps and a box trap (Fig. 1).

**Figure 1.**
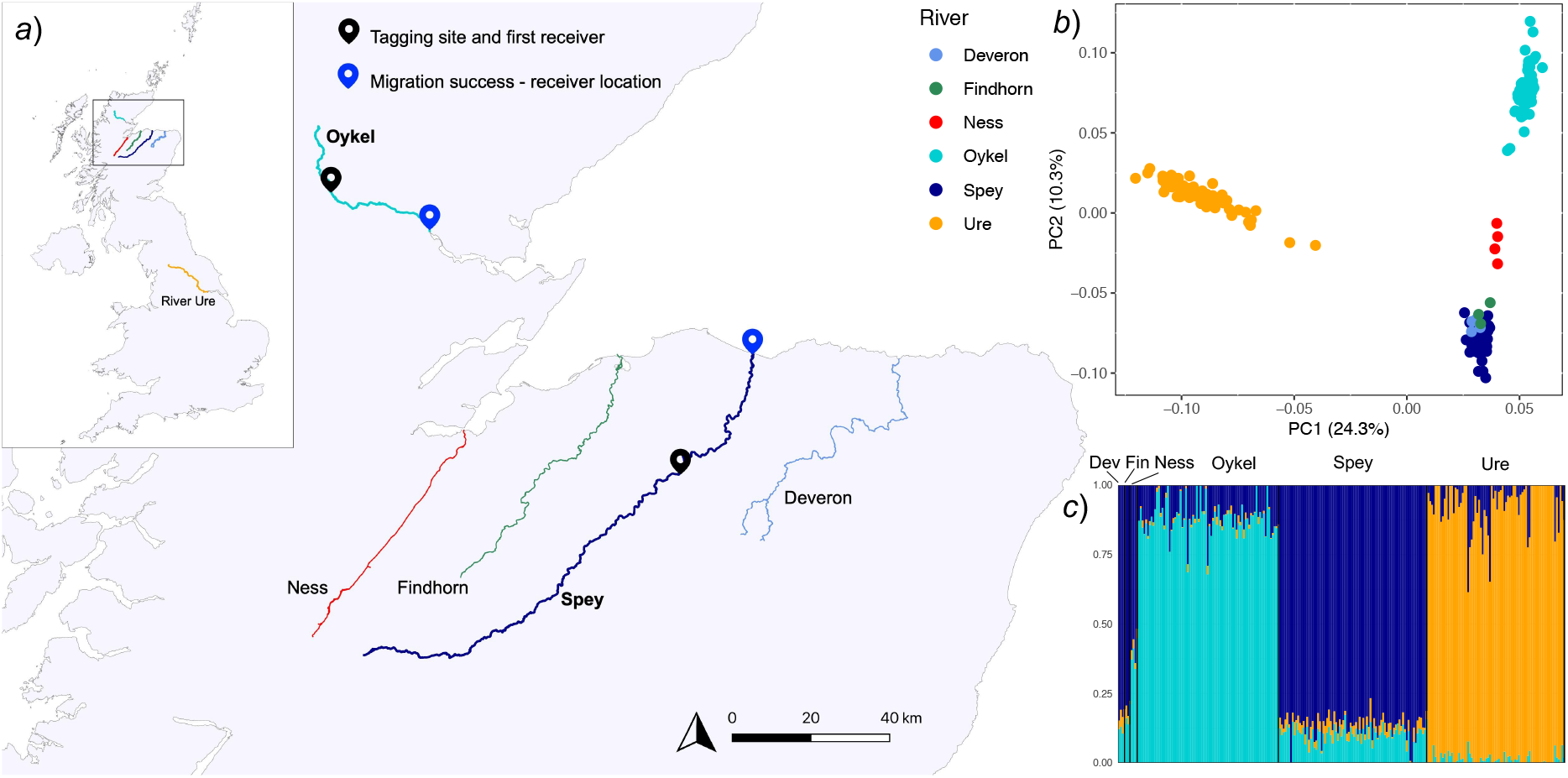
The study area (*a*) and genetic structuring (*b, c*) of salmon populations from rivers flowing into the Moray Firth (Scotland), with samples from the River Ure (England, *a*) top left panel) for comparison. Tagging locations and acoustic receivers are shown in map (*a*). The principal components analysis (PCA) and the ADMIXTURE analysis plots are based on 44,504 SNPs pruned for linkage disequilibrium. In the PCA scatterplot (*b*), dots represent individual fish, and variance (%) explained by the first and second axes are shown. Colours correspond to rivers. In the ADMIXTURE plot (K=3; *c*) each fish individual is represented by a vertical bar. ‘Dev’ and ‘Fin’ are abbreviations for the rivers Deveron and Findhorn, respectively.

Fish were anaesthetised in MS222 and tagged with Vemco V7-2L acoustic transmitters (7mm diameter, 19.5mm length, 1.5g in air, 137 dB re 1μPa @ 1m, acoustic transmission repeat cycle of 28 seconds ± 10 seconds, InnovaSea, Bedford, Nova Scotia, Canada). For more details on the tagging and release procedure see Lilly et al. (2022). Before being tagged, fish were measured (fork length, mm), weighed (g), and photographed. Photographs of the left side of each fish were taken from approximately 30 cm directly above the fish, with a Fujifilm FinePix XP130 Compact Digital Camera on a background reference scale. An adipose fin-clip was also taken from every fish and stored in 96% ethanol for later DNA extraction.

Two acoustic monitoring receivers (InnovaSea VR2Tx) were deployed in each river, one of which was immediately downstream of the tagging site (0.2 and 0.6 km in the rivers Oykel and Spey, respectively; Fig. 1). The second receiver was deployed at the river mouth (Fig. 1). Of all the salmon tagged and released in the two rivers (Oykel, n = 149, Spey n = 150), 91.9 and 96.7% respectively were detected by the first receiver after release. Of these, 78 (Oykel) and 82 (Spey) smolts were randomly selected for this study, distributed evenly between migratory outcomes ensuring a balanced design. Fish from both rivers were allocated into two groups based on their migratory outcome; 1) fish detected on the second and final river receiver were categorised as ‘successful’ river migrants, and 2) fish only detected on the first receiver were considered as ‘unsuccessful’ river migrants (Table 1, Fig. 1). To assess receiver detection efficiency, additional receivers deployed as a part of the broader telemetry study in the marine coastal waters of the Moray Firth were used. Since all smolts detected in marine waters were also detected by the two freshwater receivers, detection efficiency was determined to be 100%, meaning that no fish were wrongly miscategorised as unsuccessful migrants as a result of missed detections at the second river receiver.

**Table 1.** Classification of smolts (*n*) in the rivers Oykel and Spey based on tracking results.

| River | Unsuccessful | Successful | Total river migration distance |
| --- | --- | --- | --- |
| Oykel | 36 | 42 | 30.5 km |
| Spey | 35 | 47 | 50.1 km |

To assess whether the study rivers harboured genetically differentiated Atlantic salmon populations, the genetic variation across rivers in the Moray Firth (Fig. 1) was investigated. In addition to fish from the study rivers Oykel and Spey, Atlantic salmon smolts from the rivers Findhorn (*n* = 3; 57°25.05’ N, 3°53.35’ W), Deveron (*n* = 4; 57°30.45’ N, 2°42.35’ W) and Ness (*n* = 4; 57°27.17’ N, 4°15.35’ W) were included in the analysis. To further contextualise the relative genetic diversity of these rivers, Atlantic salmon samples from the River Ure, England (*n* = 76; 54°16.19’, N 1°44.57’ W) were also included in the analysis (Fig. 1). Fish from the Findhorn, Deveron and Ness were sampled in Spring 2019 using rotary screw traps, while fish from the Ure were captured employing backpack electric fishing equipment (Electracatch 24 V DC input, 200-400 V, 100 W, 50 Hz Pulsed DC, variable pulse width output).

### Genomic analyses

#### DNA extraction, genotyping and quality control

DNA was extracted from adipose fin samples employing a modified Mu-DNA: Tissue protocol (Sellers et al., 2018) using a solid phase reversible immobilization (SPRI) magnetic bead capture method (adapted from Rohland & Reich, 2012) to isolate high molecular weight DNA. The DNA samples were sent to the Centre for Integrative Genetics (CIGENE, Ås, Norway) for genotyping, including biological and technical replicates to ensure consistency across plates. A custom 220,000 SNP (Single Nucleotide Polymorphism) Affymetrix Axiom array designed for Atlantic salmon (see Barson et al., 2015 for details) was used for data generation. Following the manufacturer’s instructions, only SNPs categorised as PolyHighResolution and NoMinorHom were used for analyses, while SNPs with unknown position were excluded from the dataset, leaving 213,945 available loci for genomic investigation. We then performed quality control (QC) and filtering of SNPs data in PLINK version 1.9 and 2.0 (www.cog-genomics.org/plink/1.9/ and www.cog-genomics.org/plink/2.0/; Purcell et al., 2007; Chang et al., 2015). SNPs were filtered for Hardy-Weinberg equilibrium (PLINK 1.9. command: *--hwe 0*.*001*) to remove genotyping errors. Additionally, SNPs were screened for minor allele frequency (*--maf 0*.*05*) and genotype missingness (*--geno 0*.*1*), and individuals with a high rate of missing SNPs (*--mind 0*.*1*) were discarded from analyses. Full siblings were also removed using PLINK 2.0 (*--king-cutoff 0*.*25*). In the migration success analyses, these QC steps were performed separately for the rivers Oykel and Spey and resulted in the retention of 198,336 SNPs and 82 individual fish from the River Oykel and 201,475 SNPs and 78 individuals from the River Spey. The fish used in this analysis were the same employed for morphometric investigations, but included two additional individuals which were not photographed. For the regional population structure analyses (paragraph below), QC was performed on all rivers together and SNPs in high linkage disequilibrium were pruned in PLINK 1.9 (*--indep 50 5 1*.*4*) leaving 44,504 unlinked SNPs available for analysis.

#### Regional rivers genetic structuring

To investigate the genetic variation across rivers in the Moray Firth (Fig. 1), a principal component analysis (PCA) was performed in PLINK 1.9 using individuals from the rivers Oykel, Spey, Findhorn, Deveron, Ness and Ure. Using the same samples, ADMIXTURE v.1.3. (Alexander, Novembre & Lange, 2009) was used to infer the most likely number of genetic clusters (K, testing from K=1 to K=6), that was determined based on the lowest cross-validation error.

#### Outlier analysis and gene annotation

To detect SNP markers with unusually high levels of allelic differentiation between successful and unsuccessful migrants in each river, two different approaches were computed using the unpruned SNP dataset. In the first approach, the R (R Core Team, 2022) package ‘OutFLANK’ v. 0.2 (Whitlock & Lotterhos, 2015), which estimates among-groups Fst for each locus was used. For the second, the allele-based chi-squared association test in PLINK 1.9 (command: --assoc) was implemented. See code (https://tinyurl.com/salmon-migration-success) for details about parameters used to run these analyses. Outlier loci in ‘OutFLANK’ were identified by applying a q-value < 0.05 threshold, while outliers in the association test in PLINK were determined as the top 0.1% SNPs ranked by P-values. Acknowledging the inherent risk of false positives in genome scan analyses (Luu, Bazin & Blum, 2017), a robust bootstrapping methodology was employed. For each river, 200 bootstrap replicate datasets were generated by randomly removing one fish from each of the migratory groups (successful and unsuccessful). Each of these datasets was examined independently, with 100 analysed using ‘OutFLANK’ and the other 100 using the PLINK association test. Only the outliers that were consistently detected in all 200 bootstrap replicates by both methods were retained for subsequent analysis. Outliers were visualised using the ‘qqman’ v. 0.1.8 R package (Turner, 2014). A PCA of the outliers was computed in PLINK on the complete dataset including all fish and the resulting PCA plot was employed to visually test if these outlier SNPs effectively separated successful and unsuccessful migrating salmon smolts in the two rives.

For each river, the ten genes closest to each outlier SNP were extracted using the *closest* function in the software bedtools 2.29.1 (Quinlan & Hall, 2010) and the genes within 10 kb upstream and downstream of outlier SNPs were filtered in R (Wellband et al., 2019). The potential functions of these genes were assessed by examining the gene ontology (GO) biological process terms associated with each gene, using the R package ‘Ssa.RefSeq.db’ v.

1.2 (Grammes, 2016) and literature searches. For these analyses, the NCBI Salmo salar Annotation Release 100 (ICSASG_v2) was used as a reference genome.

### Morphometric analyses

#### Landmark digitisation

Fish morphology was analysed using length (mm), weight (g), Fulton’s condition factor (K; Nash, Valencia & Geffen, 2006) and geometric morphometrics (GM). The GM analyses were based on photographs of 158 salmon (Spey *n* = 77, Oykel *n* =81). The images of each fish were imported into tpsUtil v. 1.78 (Rohlf, 2019) and randomly shuffled using the *Randomly order specimens* function so that the operator was blind to the river-of-origin of the specimens. Nine fixed and 4 semi-landmarks (Fig. 2) were digitised on each image by a single operator using tpsDig v. 2.31 (Rohlf, 2017) using a subset of landmarks from the scheme proposed by Moccetti et al. (2023). Furthermore, five linear body measurements (Fig. 2) used as proxy of body slenderness which has been associated to swimming ability were also included (e.g., Pakkasmaa & Piironen, 2001; Drinan et al., 2012). Landmark coordinates were imported into R and analysed using the ‘geomorph’ and ‘RRPP’ v. 4.0.4 packages (Adams et al., 2021; Baken et al., 2021; Collyer & Adams, 2021). Preliminary analysis revealed body bending as a major source of shape variation in the dataset. This was corrected by employing landmarks 1, 14, 15 and 11 (see Moccetti et al., 2023 for details). All subsequent analyses were performed on landmarks 1-13 only. PCA plots were produced with the ‘ggplot2’ package (Wickham, 2016).

**Figure 2.**
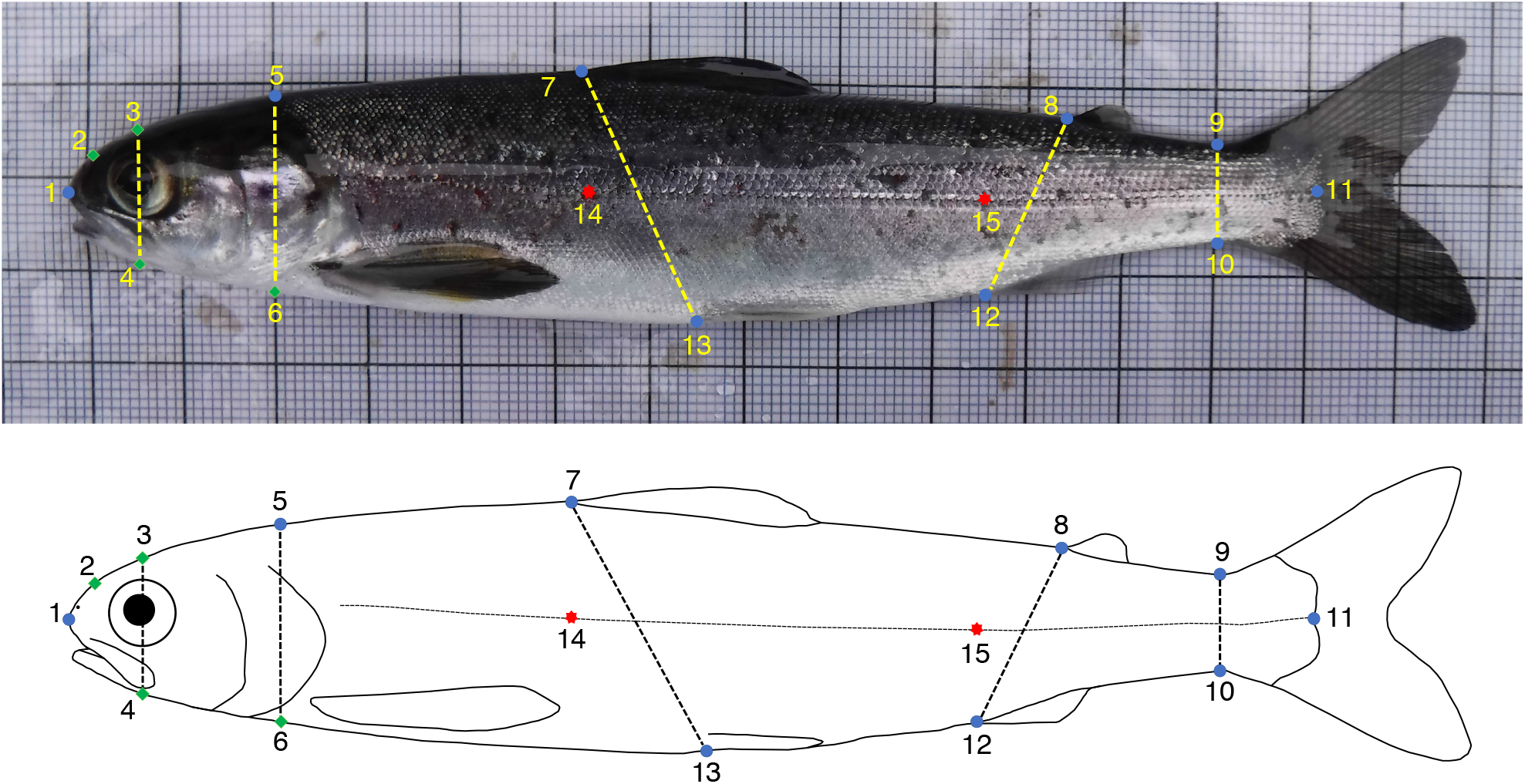
Fixed (blue circles) and semi (green diamonds) landmarks and linear measurements used for the geometric morphometrics analyses of Atlantic salmon smolts (image modified from Moccetti et al, 2023). Landmarks 14 and 15 (red stars) were used to correct for body arching. **(1)** Tip of snout; **(2)** Midpoint between 1 and 3; **(3)** Directly above middle of eye; **(4)** Perpendicular to 3; **(5)** Dorsal surface posterior of cranium; **(6)** Perpendicular to 11; **(7)** Anterior insertion point of dorsal fin; **(8)** Anterior insertion point of adipose fin; **(9)** Dorsal insertion point of caudal fin; **(10)** Versal insertion point of caudal fin; **(11)** Posterior midpoint of hypural plate; **(12)** Anterior insertion point of anal fin; **(13)** Anterior insertion point of ventral fin; **(14)** Lateral line - perpendicular to 7; **(15)** Lateral line - perpendicular to 12.

The following analyses were performed separately for each river. First, we tested whether successful and unsuccessful migrating fish in the two rivers were different in length (mm), weight (g) or Fulton’s condition factor using *t*- and *Mann-Whitney U*-tests depending on the data distribution. Fulton’s condition factor (K) was calculated as: *K = 100 x W*.*L*^*-3*^, where *W* = weight (g) and *L* = length (cm). We next tested for a difference in body shape between successful and unsuccessful fish, and whether these differences were consistent across rivers. A generalised Procrustes analysis (GPA) was performed to remove effects not related to body shape through translation, scaling and rotation of the landmark configurations (Rohlf & Slice, 1990). The residual effect of fish size on body shape was tested using Procrustes ANOVAs, with Procrustes coordinates used as a response variable and log centroid size used as an independent variable with a randomised residual permutation procedure (10,000 iterations). No significant effect of size on shape was found in either river (*p* > 0.05). To visualise body shape of successful and unsuccessful fish, a PCA was performed on the Procrustes-aligned coordinates of fish of each migration category from each river. Procrustes ANOVAs and *t*-tests were subsequently used to test for differences in body shape and linear distances between fish with different migratory outcomes. For linear distance and length, weight and condition factor comparisons, significance values were Bonferroni corrected to limit the increased error rate correlated with multiple testing (Rice, 1989).

## Results

### Genomic analyses

#### Regional rivers genetic structuring

The study rivers Spey and Oykel were genetically differentiated from one another (Fig. 1). The second Principal Component (PC2) successfully separated the fish from the River Ness, geographically located between the Oykel and the Spey, from all the other rivers (Fig. 1). Fish from the rivers Deveron and Findhorn clustered with the spatially adjacent River Spey (Fig. 1). PC1 separated the more geographically distant River Ure, situated in northern England, from all Scottish rivers flowing into the Moray Firth (Fig. 1). Similarly, ADMIXTURE analysis identified three different genetic clusters consisting of the rivers Oykel, Spey and Ure, with the rivers Deveron, Findhorn clustering with the Spey, while the Ness was admixed with the rivers Oykel and Spey.

#### Genomic regions linked to migratory outcome

There was a consistently high false discovery rate (FDR) observed across methods and bootstrap replicates in both rivers. On average, only 11.7% of outlier SNPs were detected by ‘OutFLANK’ or PLINK in all 100 replicates. Specifically, the FDR was 10.8% (‘OutFLANK’) and 10.7% (PLINK) in the Oykel dataset and 14.4% (‘OutFLANK’) and 10.7% (PLINK) in the Spey. Seventy outlier SNPs were consistently detected by both methods in all bootstrap replicates for migrating fish in the River Oykel, and 67 outliers for fish from the River Spey (Supplementary files 1, 2). None of the outlier SNPs were found in fish from both rivers. The PCA computed on this subset of outlier loci confirmed their ability to distinguish between successful and unsuccessful fish along PC1 axis for each river (Fig. 3).

**Figure 3.**
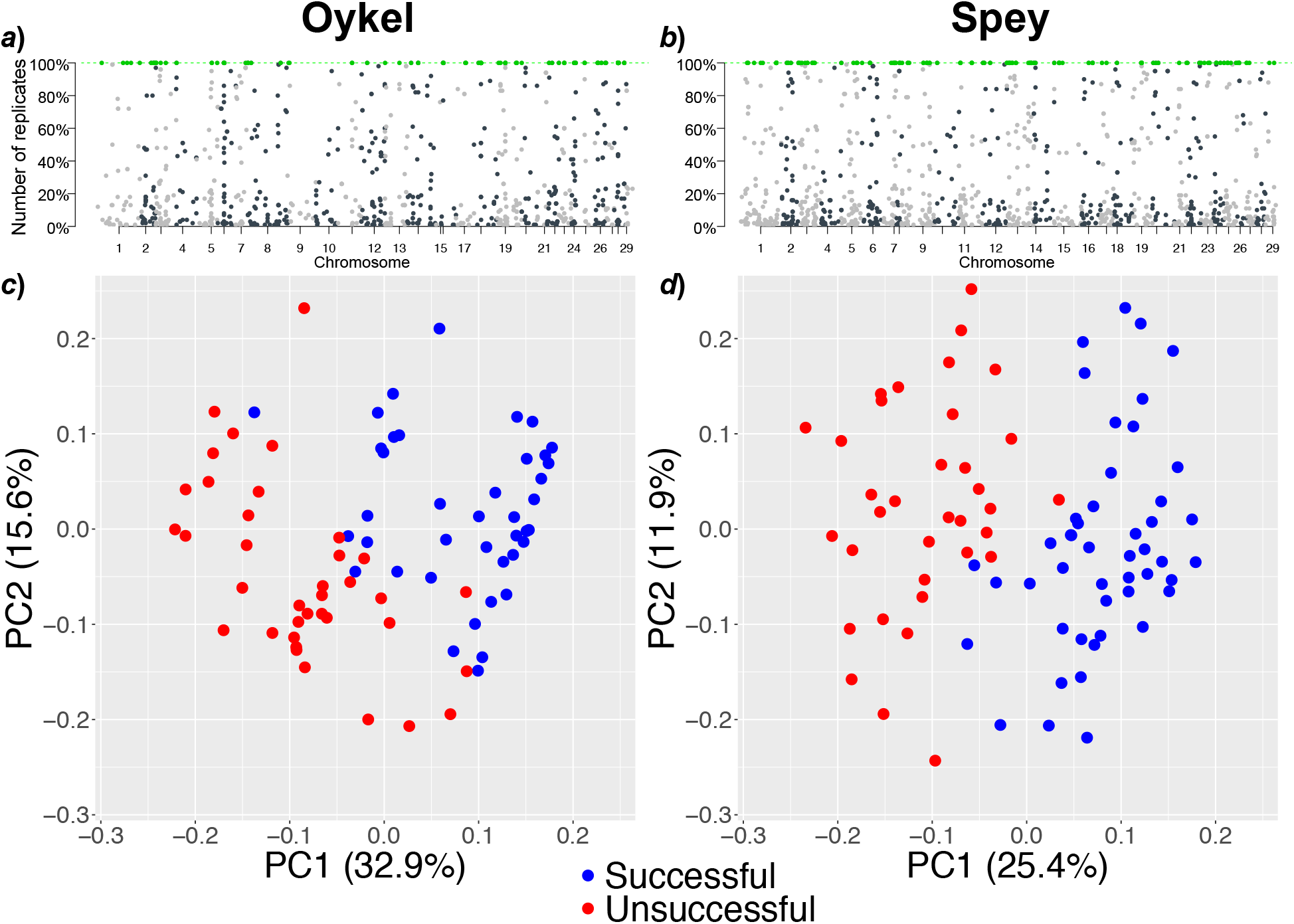
The Manhattan-style plots (*a, b*) show all outlier SNPs (dots) identified in bootstrap replicated datasets using ‘OutFLANK’ in each river. The outliers consistently detected in 100% of replicates and used for analysis are highlighted in green. The y-axis shows the proportion of replicated datasets where each individual outlier SNP was identified. The x-axis displays the position of the SNPs along the genome with chromosome numbers. The analogous plots for the association test in PLINK are shown in Supplementary Figure 1. *c* and *d*; Principal components analysis scatterplots based on 70 (Oykel) and 67 (Spey) outlier SNPs between successful (blue) and unsuccessful (red) migrant Atlantic salmon smolts. Each dot represents an individual fish. Variance (%) explained by the first and second axes is also shown.

#### Gene annotation

There were 50 and 48 putative coding regions (hereafter genes) within 10 kb of the outlier SNPs’ locations in the Oykel and Spey fish, respectively (Supplementary file 3). None of these genes were identified as outliers in fish from both rivers. Eight and 12 genes contained more than one outlier SNP within the 10kb region in the Oykel and Spey samples, respectively (Supplementary file 3). The two genes enclosing the highest number of outlier SNPs were the *anion exchange protein 2-like* (encompassing 18 SNPs, River Oykel) and the *collagen alpha-1(I) chain-like* (encompassing 10 SNPs, River Spey).

### Morphological differences between successful and unsuccessful salmon

There were no differences in length, weight or Fulton’s condition factor between successful and unsuccessful migrating smolts (*p* > 0.07; Supplementary Figure 2, Supplementary Tables 1, 2). Procrustes ANOVAs based on the 13 landmark coordinates did not show significant differences in body shape between successful and unsuccessful smolts in either of the two rivers (*p* > 0.31; Fig. 4, Supplementary Table 3). Likewise, after Bonferroni correction (new *alpha* value = 0.005), comparison of body linear measurements did not show any significant difference between migrating groups (*p* > 0.04; Supplementary Tables 4, 5).

**Figure 4.**
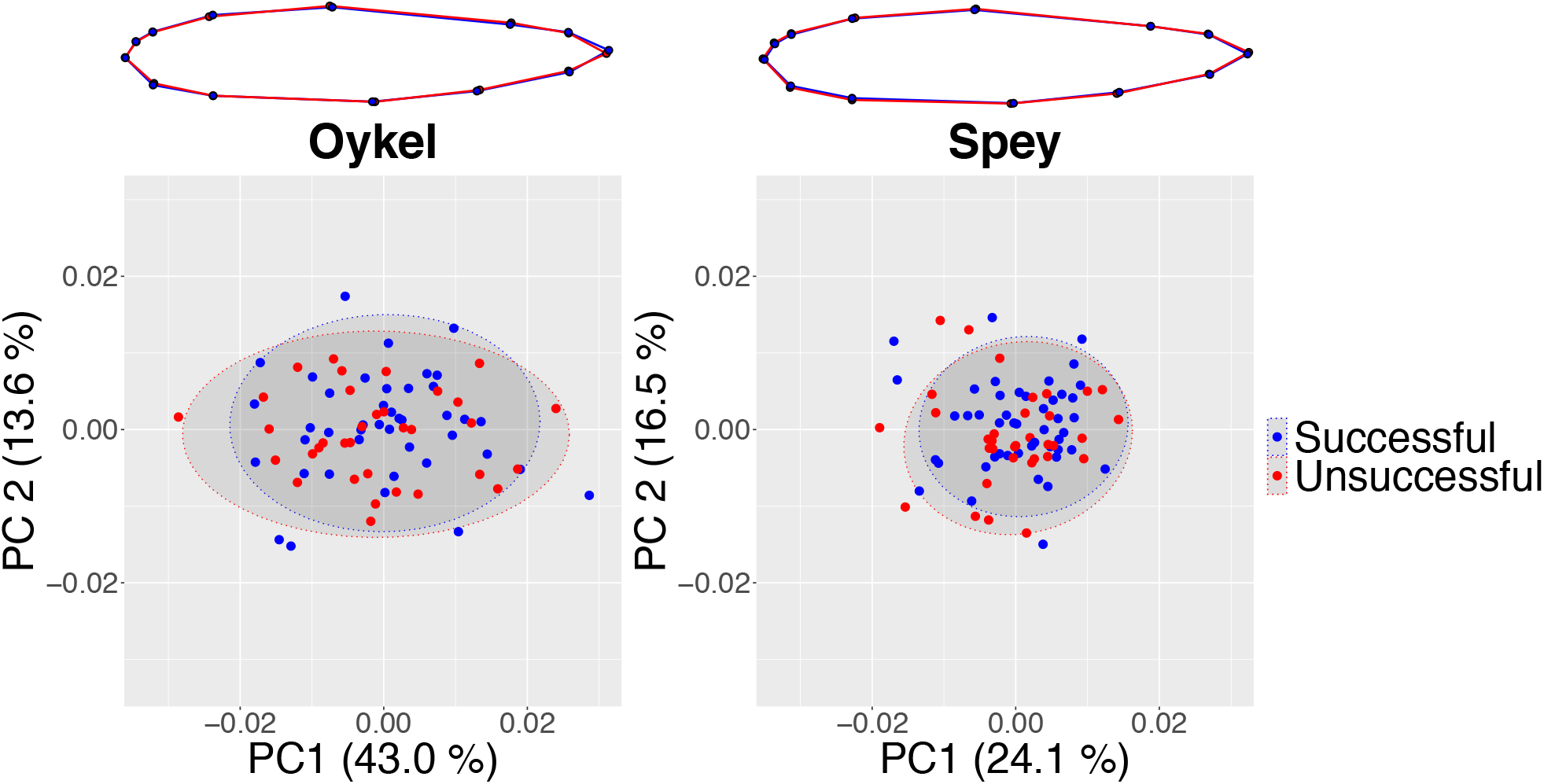
Mean body shape projections (top) and principal components analysis scatterplots (lower panel) show an absence of shape difference between the migratory groups. Procrustes-aligned coordinates of successful (blue) and unsuccessful (red) Atlantic salmon, where dots represent individual fish are shown below. Variance (%) explained by the first and second axes and 95% confidence ellipses are displayed. Projections show a complete overlap of the blue (successful) and red (unsuccessful) lines in both rivers despite magnifying morphological differences three times to aid visualisation.

## Discussion

Our work shows that distinct SNP sets were significantly differentiated between Atlantic salmon smolts making successful migrations to sea and those that failed to migrate to sea in two different rivers. In both rivers, the outlier SNPs predicting individual migration success were near several genes that could be relevant for migration, but we found no evidence of phenotypic differences in body shape between successful and unsuccessful Atlantic salmon river migrants.

Categorising genes containing outlier SNPs by biological function highlighted similar processes across the study rivers. In fish from both rivers, genes putatively linked to osmoregulation, immunity, stress, and nervous, sensory, muscular, skeletal and cardiovascular system development and activity were detected.

Candidate genes linked to general neuronal, cardiovascular and skeletal functions may play an important role in migration, but a direct link to smolt migration success is hard to determine. Furthermore, given the susceptibility of gene annotation to false positives, it is important to exercise caution when attempting to establish such correlations (Pavlidis et al., 2012). Nevertheless, osmoregulation and immune response are processes shown to play an important part in salmonid migration. In the Oykel and in the Spey outlier SNPs were located within or near (within 10 kb) several candidate osmoregulatory genes. These genes were associated with a range of processes including ion transmembrane and water transport, renal activity, response to salt stress and hyperosmotic response. Noteworthy it is the identification of the *anion exchange protein 2-like* gene, encompassing 18 of the 70 outlier SNPs detected in the Oykel. This gene is associated to GO terms involved in osmoregulatory processes, such as chloride and bicarbonate transmembrane transport (Wilson, Wilson & Grosell, 2002; Grosell, 2006). Osmoregulation and individual ability to undergo physiological changes required for seawater entry has been shown to be important to increase chances of survival and predator avoidance in seaward migrating salmonid smolts (Kennedy, Gale & Ostrand, 2007) and could play a role in migration success of Atlantic salmon smolts in the last tidal kilometres of the Oykel and Spey where transitional zone between freshwater and saltwater occurs.

Immunity related and stress response genes were also widely detected in association with the outlier SNPs separating successful and unsuccessful river migrants in fish from both the Oykel and Spey. Studies using proteomics in Pacific salmon (*Oncorhynchus* spp.) have found significant correlation between migratory outcome, expression of specific immune-related genes and viral and parasite-induced infection burden (Miller et al., 2011; Jeffries et al., 2014; Furey et al., 2021; Mauduit et al., 2022). The stress hormone cortisol has also been found to be a good predictor of migration success in salmonids (Birnie-Gauvin et al., 2019). Our findings now highlight the potentially important role of pathogens driven selection in Atlantic salmon migration success. An additional factor that requires further investigation, is the possibility that there are individual differences in response to the tagging process, since there are also immune genes annotated with GO terms involved in blood coagulation and response to wounding. To determine migration patterns, all the fish in our study were tagged, so although this is not a confounding factor in our design, this finding warrants further investigation.

While particular SNP sets allow us to predict migratory outcome of Atlantic salmon smolts in the Oykel and Spey, analyses of length, weight, body condition and body shape did not find any significant difference between successful and unsuccessful migrants in either of the rivers. This is somewhat surprising given the importance that morphology plays in swimming performance in fish (Webb, 1978; Webb, 1984; Fisher & Hogan, 2007; Langerhans & Reznick, 2010; Assumpção et al., 2012), the specific hydrodynamic characteristics required to effectively migrate in running waters (Langerhans & Reznick, 2010; Brodersen et al., 2014), and that size, body shape and condition may also be important in anti-predatory behaviour (Domenici et al., 2007). However, we did find a difference in smolt morphology between rivers (Moccetti et al., 2023) and there is genetic differentiation of the Oykel and Spey salmon populations indicating reproductive isolation between these geographically close populations, likely facilitated by fine-scale homing. Different evolutionary and demographic histories, (evidenced by the geographic structure we find between river populations) combined with different contemporary ecological selection pressures will therefore lead to different traits being linked to migration success. For example, there were no genes that contained outlier SNPs nor GO processes in common in Atlantic salmon from both the Oykel and Spey.

Overall, we found that migratory outcome for individual salmon smolts in given rivers, in a given year, could be predicted from a subset of SNPs consistently detected through bootstrapping approach. We next need to understand the ecological and environmental factors which could determine those subsets, by adding temporal replication so that we can better understand the limits of our study. From an evolutionary and conservation point of view, the mechanism through which the observed genetic diversity could be maintained needs to be identified, given that migration failure should theoretically quickly purge polymorphism at selected SNPs from a population. We propose that variation in life history could maintain standing genetic variation for environmentally driven balancing selection (Mérot et al., 2020). ‘Partial’ migration (Shaw, 2016), where only a portion of the population migrates, is common in several taxa and may be responsible for maintaining high genetic diversity if migratory and resident individuals interbreed (Pulido, 2011). Within-population differences in migratory strategies (e.g. timing, duration and routes) between age-classes and sexes are also well known phenomena (Cristol, Baker & Carbone, 1999). These sub-groups are exposed to different biotic and abiotic conditions potentially selecting for different genotypes that may maintain the gene pool diversity within the population (Dingle & Drake, 2007; Wittmann et al., 2017; Briedis & Bauer, 2018). Alternative migratory and reproductive tactics are well documented in Atlantic salmon (Fleming, 1998; Thorstad et al., 2010; Birnie-Gauvin, Thorstad & Aarestrup, 2019), thus individuals with different life histories experiencing temporally and spatially fluctuating selection can interbreed and induce genetic mixing. Typically, the life history of Atlantic salmon involves seaward migration followed by a return to their natal river to spawn, but a number of males (and occasionally females, Birnie-Gauvin, Thorstad & Aarestrup, 2019) become sexually reproductive in freshwater as morphological juveniles before migrating to sea (‘precocious male parr’; Lepais et al., 2017). Their contribution to paternity could be substantial (ca. 60% in one study, Saura et al., 2008). The number of years spent in freshwater before smolting, and at sea before upstream spawning migration can also vary considerably (Thorstad et al., 2010). Finally, unlike Pacific salmon species, a non-negligible proportion of Atlantic salmon survive reproduction (especially females), return to the ocean as ‘kelt’ and spawn multiple times (Hedger et al., 2009). Weather and ecological conditions can change dramatically among and within years inducing different selective pressures on migrating smolts and other salmon life stages. For instance, variations in water discharge and temperature may affect ecological factors such as migration timing (Thorstad et al., 2010), predation (Kennedy, Gale & Ostrand, 2007; Hostetter et al., 2012) and pathogen infection (Wagner et al., 2005) as well as passage of artificial migration barriers (Marschall et al., 2011). Clearly, all these variables may differentially alter the allele frequencies under selection and help maintain standing genetic variation. Straying between rivers could also be a source of genetic diversity (Palstra, O’Connell & Ruzzante, 2007; Keefer & Caudill, 2014), although we found no evidence of this in our study.

From a conservation point of view, understanding and predicting these selection pressures could be invaluable in managing existing populations, and could inform stock selection where hatchery-reared individuals are used to augment populations (Jepsen, Nielsen & Deacon, 2003; Koed et al., 2020; Waples, Naish & Primmer, 2020).

Overall, our findings show that migration success could be linked to specific genotypes and highlight the importance of preserving genetic diversity for conservation, to allow populations to respond to potential heterogeneity between years, and the increased variability that long-term climate change may produce. Our next challenge is to understand in detail the selection pressures and associated genetic changes in populations facilitating conservation success and ensuring a future for these iconic species.

## Acknowledgments

We would like to thank the Spey Fisheries Board and the Kyle of Sutherland Fisheries Trust for their help and support with this project. We would also like to thank Jonathan Archer, Georgios Kyriakou, Fraser Brydon, Jessica Whitney, Mustafa Soganci and Rowan Smith for their help collecting the data for this study.

## Ethics

For fish samples from the rivers Oykel, Spey, Deveron, Findhorn and Ness, the care and use of experimental animals complied with the UK Home Office animal welfare laws, guidelines and policies (UK Home Office Licence PPL 70/8794) and was approved by the University of Glasgow Animal Welfare and Ethics Review Board (AWERB). Atlantic salmon from the River Ure were treated in compliance with the UK ASPA (1986) Home Office project licence number PD6C17B56.

