## Supplementary Figures and Tables for "Individual genotype but not phenotype predicts river migration success in Atlantic salmon"

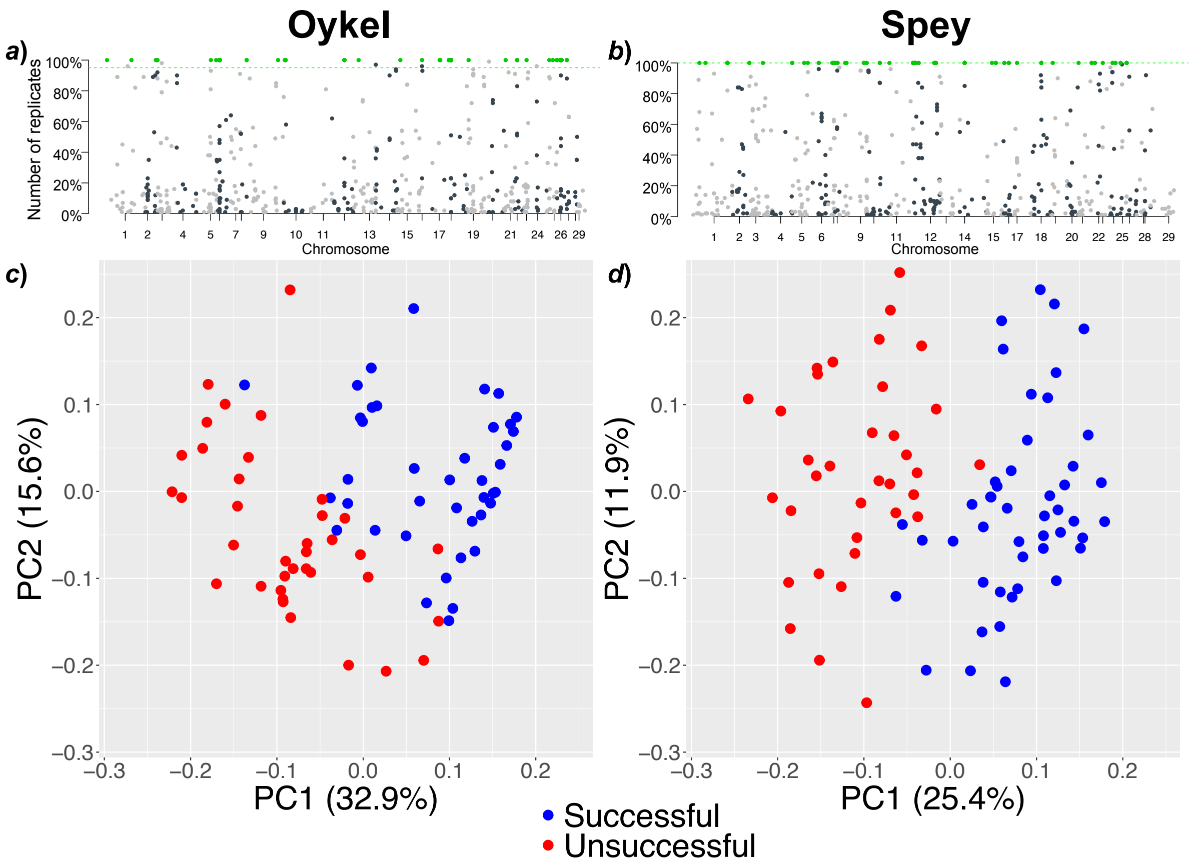


**Supplementary Figure 1.** The Manhattan-style plots (*a*, *b*) show all outlier SNPs (dots) identified in bootstrap replicated datasets using the allele-based chi-squared association test in PLINK in each river. The outliers consistently detected in 100% of replicates and used for analysis are highlighted in green. The y-axis shows the proportion of replicated datasets where each individual outlier SNP was identified. The x-axis displays the position of the SNPs along the genome with chromosome numbers. The analogous plots for the association test in 'OutFLANK' are shown in Figure 3 in the main text. *c* and *d*; Principal components analysis scatterplots based on 70 (Oykel) and 67 (Spey) outlier SNPs between successful (blue) and unsuccessful (red) migrant Atlantic salmon smolts. Each dot represents an individual fish. Variance (%) explained by the first and second axes is also shown.

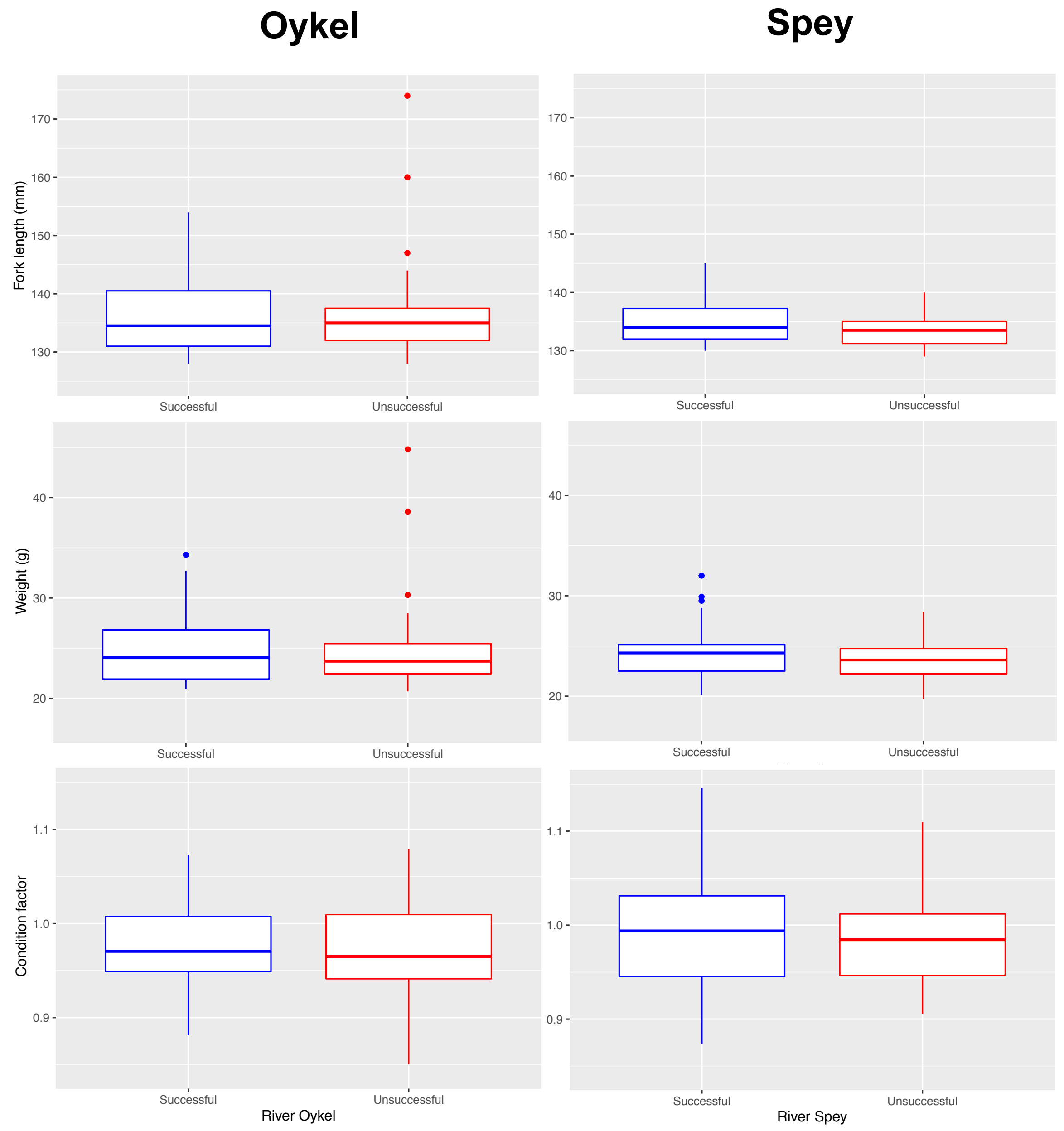


**Supplementary Figure 2.** Boxplots of length (mm), weight (g) and Fulton's condition factor to compare successful and unsuccessful Atlantic salmon smolts in the rivers Oykel and Spey.

**Supplementary Table 1.** Summary statistics of body metrics comparisons between successful and unsuccessful Atlantic salmon smolts in the River Oykel. *M-W U* = *Mann-Whitney U*- test.

| **Metric** | **Test** | **Df** | ***t*** | ***W*** | ***P*-value** |
|---|---|---|---|---|---|
| Length | *t*-test | 55.228 | -0.51252 | - | 0.6103 |
| Weight | *M-W U* | - | - | 774 | 0.6936 |
| Condition factor | *t*-test | 74.479 | 1.0039 | - | 0.3187 |

**Supplementary Table 2.** Summary statistics of body metrics comparisons between successful and unsuccessful Atlantic salmon smolts in the River Spey. *M-W U = Mann-Whitney U*- test.

| **Metric** | **Test** | **Df** | ***t*** | ***W*** | ***P*-value** |
|---|---|---|---|---|---|
| Length | *M-W U* | - | - | 951 | 0.1452 |
| Weight | *t*-test | 78.108 | 1.8104 | 774 | 0.07408 |
| Condition factor | *M-W U* | - | - | 841 | 0.6913 |

**Supplementary Table 3.** Procrustes ANOVA summary statistics of effect of migratory outcome on the body shape of Atlantic salmon smolts in the rivers Oykel and Spey.

| **River** | **Df** | **SS** | $r^2$ | ***F*** | ***Z*** | ***P*-value** |
|---|---|---|---|---|---|---|
| Oykel | 1 | 0.0003318 | 0.0146 | 1.1109 | 0.47321 | 0.3132 |
| Spey | 1 | 0.0002305 | 0.01351 | 1.0817 | 0.35458 | 0.3659 |

**Supplementary Table 4.** Summary statistics of *t*-tests comparing body linear measurements between successful and unsuccessful Atlantic salmon smolts in the River Oykel.

| **Linear distance (landmark numbers)** | **Df** | ***t*** | ***P*-value** |
|---|---|---|---|
| 3-4 | 72.723 | 0.82542 | 0.4118 |
| 5-6 | 71.748 | 0.81866 | 0.4157 |
| 7-13 | 74.025 | -1.6729 | 0.0986 |
| 8-12 | 62.254 | 0.13079 | 0.8964 |
| 9-10 | 71.747 | 1.9853 | 0.0509 |

**Supplementary Table 5.** Summary statistics of *t*-tests comparing body linear measurements between successful and unsuccessful Atlantic salmon smolts in the River Spey.

| **Linear distance (landmark numbers)** | **Df** | ***t*** | ***P*-value** |
|---|---|---|---|
| 3-4 | 68.916 | -1.5172 | 0.1338 |
| 5-6 | 73.741 | -2.1179 | 0.0376 |
| 7-13 | 77.503 | 0.16284 | 0.8711 |
| 8-12 | 76.554 | -0.93755 | 0.3514 |
| 9-10 | 73.045 | -0.21804 | 0.8280 |
